# Soil viral communities are structured by pH at local and global scales

**DOI:** 10.1101/2021.10.20.465127

**Authors:** Sungeun Lee, Jackson W. Sorensen, Robin L. Walker, Joanne B. Emerson, Graeme W. Nicol, Christina Hazard

## Abstract

Viruses shape microbial community structures, impacting metabolic pathways and influencing biogeochemical cycles. Despite their importance, the influence of biotic and abiotic factors on viral community structures across environmental gradients in soil is relatively unknown compared to their prokaryotic hosts. While soil pH strongly influences microbial community structure, it is unclear whether there is a similar influence on soil virus communities. In this study, prokaryotic and viral communities were characterized in soils sampled from the extremes of a long-term pH-manipulated soil gradient (pH 4.5 and 7.5), and viral populations were compared to those in a variety of soil ecosystems ranging in pH (4.0 – 7.5). Prokaryotic and viral community structure were significantly influenced by soil pH at the local scale. Of 1,910 viral operational taxonomic units (vOTUs), 99% were restricted to pH 4.5 or 7.5 soil only. These were compared in gene sharing networks of populations from six other European and North American soil systems. A selection of viral clusters from acidic and neutral pH soils were more associated with those from the local gradient pH 4.5 or 7.5 soils, respectively. Results indicate that as with prokaryotes, soil pH is a factor structuring viral communities at the local and global scale.

## Main

Viruses play a major role in controlling the abundance, structure and evolution of microbial communities through cell lysis and the release of nutrients and modulation of host cell metabolism during infection [1]. In soils, viruses are diverse [2] and abundant [3], likely infect the majority of bacteria and archaea, and have considerable potential to impact carbon sequestration, nutrient cycling and other ecosystem functions [2,4,5]. Amongst the many knowledge gaps in soil viral ecology includes a basic understanding of the biotic and abiotic drivers of viral communities at both the local and global scale.

Host communities are likely the strongest factor for defining viral community structures with virus populations derived from infection of susceptible hosts. As with other environments, soil bacterial and viral community dynamics co-vary, with the susceptibility of hosts to infection from individual viruses varying over time [6] or viral community shifts occurring as an indirect result of nutrient input altering host community structure [7]. While the presence of specific hosts will influence the composition of virus communities, the host range of viruses may also play a role in defining whether their community dynamics vary to the same extent as prokaryotic communities over physicochemical gradients in soil [8]. Closely related host populations (e.g. of the same genus) may be adapted to growth under different conditions, resulting in niche differentiation and contrasting distribution across an ecological gradient (e.g. pH), but it is unclear whether narrow (e.g. single population) or broad (genus to phylum) host ranges of their associated viruses reduce the relative variation in virus community structure compared to prokaryotes. In addition, changes in soil physicochemical characteristics such as pH, temperature and moisture, may also directly impact the physical integrity and dispersal of viruses [9]. As soil pH is a major determinant of prokaryotic community structures at local and global scales [10,11,12], it is possible that virus communities may also exhibit pH-influenced community structures at similar scales.

To examine the effect of soil pH on double-stranded DNA virus community structure at a local scale, triplicate soil samples were taken from pH 4.5 and 7.5 subplots of a well-characterized soil pH gradient. At this site, prokaryotic community structure has been shown to vary in response to pH [13,14], and distinct virus populations infecting methylotrophic communities associated with contrasting soil pH have recently been observed [15]. Both non-targeted total community metagenome and virus-targeted virome libraries were prepared as previously described [15] for comparing prokaryotic and viral community structures. From all six soil samples, 908 and 763 million quality-filtered metagenome and virome reads were obtained, respectively, with 9,928 metagenome and 13,533 virome contigs ≥10 kb produced after assembly. Of these, 105 and 7,684 contigs were predicted to be of viral origin from metagenomes and viromes, respectively, and clustered into 1,910 viral operational taxonomic units (vOTUs) (Supplementary Methods; Table S1). For characterizing prokaryotic community structures, reads were annotated and prokaryotic 16S rRNA gene fragments were extracted and classified into 2,312 OTUs (Supplementary Methods). Sequencing depth of each sample and sampling size of pH 4.5 and 7.5 soil was sufficient to capture vOTU richness, although further sequencing and sampling may have increased the number of 16S rRNA OTUs recovered (Fig. S1).

Of the metagenomic reads, 22.7% were taxonomically defined, with Actinobacteria (pH 4.5, 48 ± 1.5% (standard deviation); pH 7.5, 39 ± 0.5%) and Proteobacteria (pH 4.5, 27 ± 0.5%; pH 7.5, 33 ± 0.2%) dominating in both pH soils (Fig. 1a). Similar to other soil viral studies, only a small proportion of the prokaryotic viral community was taxonomically defined. Using a gene-sharing network [16], 1.4% of viral contigs (7.0% of vOTUs) clustered with RefSeq genomes of Siphoviridae, Podoviridae and Myoviridae (Fig. 1a). Prokaryote hosts associated with viral contigs were predicted using both gene-sharing network and host-virus shared gene homology [17], linking a total of 24.5% of viral contigs (29.1% of vOTUs) (Table S2). Consistent with the dominant prokaryotic taxa, the majority of predicted viral hosts for both pH soils were Actinobacteria (pH 4.5, 76.9 ± 0.003%; pH 7.5, 64.6 ± 0.002%) and Proteobacteria (pH 4.5, 18.5 ± 0.003%; pH 7.5, 30.2 ± 0.001%) (Fig. 1a and Table S2).

**Fig. 1:**
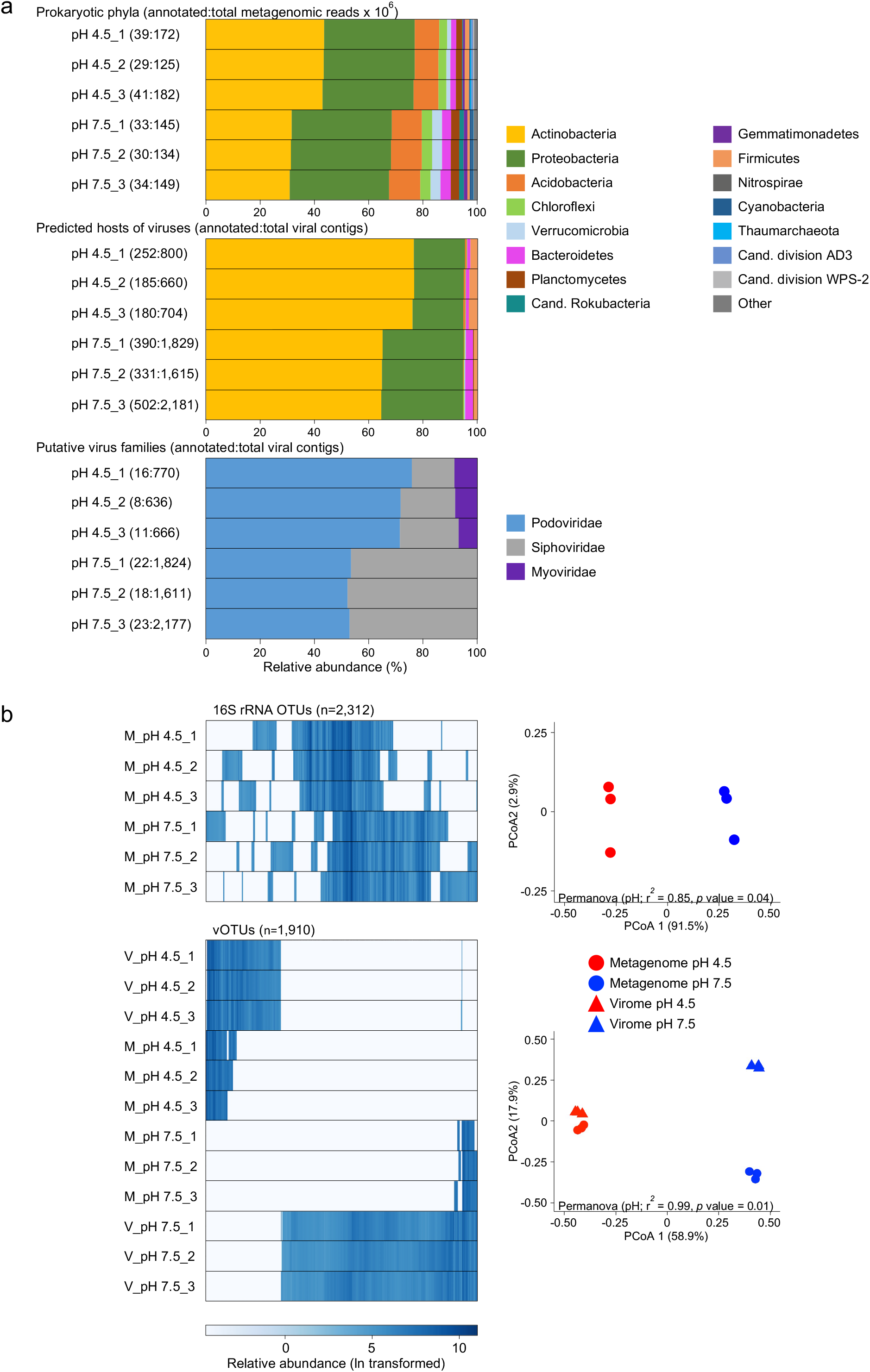
Taxonomic composition and community structure of prokaryotic and virus communities in pH 4.5 and 7.5 replicate soil samples taken from the ends of a contiguous pH gradient. a) Relative abundance of taxonomically-defined prokaryotes, taxonomically-defined viruses and the predicted hosts of viruses. For prokaryotes, reads from metagenomes were annotated at the phylum level using the NCBI nr database. Numbers in parenthesis denote the number of mapped reads:total reads analyzed. Viral contigs ≥10 kb were taxonomically defined at the family level based on gene-sharing network analysis. Host prediction of viruses was determined by gene-sharing network and gene homology analyses. Numbers in parenthesis denote the number of annotated:total reads or contigs for each sample, and plots display the relative proportion of annotated reads only (i.e. annotated reads of prokaryotes or reads mapped to annotated viral contigs). b) Normalized relative abundance of individual 16S rRNA OTUs and vOTUs in soil samples determined by read-mapping. Only vOTUs where reads were mapped with ≥1x coverage over 75% contig breadth were included. Ordinations show the principal coordinate analysis of Bray-Curtis dissimilarities derived from relative abundance tables. For virus communities, reads from both viromes (V) and metagenomes (M) were analyzed for each sample. Details of all methods used are provided in Supplementary Information.

A high resolution analysis of the distribution of individual 16S rRNA OTUs and vOTUs demonstrated distinct structures between pH 4.5. and 7.5 soils for both prokaryote and viral communities (Fig. 1b). Specifically, 38.6% of OTUs (pH 4.5 OTUs, 263; pH 7.5 OTUs, 630) and 99.0% of the vOTUs (pH 4.5 vOTUs, 524; pH 7.5 vOTUs, 1,361) were found in only one soil pH. While no significant differences in prokaryote alpha-diversity (Shannon, Simpson and richness) were observed, viral (virome) alpha-diversity was significantly greater in pH 7.5 compared to pH 4.5 soil (Student’s t-test, *p* value <0.05) (Table S3). As absorption of viruses to soil organic particles may decrease with increasing pH [18], this could have contributed to the observed differences in alpha-diversity despite a neutral pH buffer being used for extracting viral particles in both soils. Interestingly, the relative proportions of virus-encoded putative auxiliary metabolic genes (AMGs) identified in vOTUs were similar in both pH soils, indicating that pH did not select for distinct AMG composition across this pH gradient (Fig. S2).

While viromes produced 73× more assembled viral contigs than metagenomes, read mapping of individual virome and metagenome reads to all vOTUs demonstrated that both approaches produced distinct viral community profiles between soils (Fig. 1b). Interestingly, decreasing the breadth (length of contig covered by mapped reads) threshold (<75%) for defining vOTU detection disproportionately increased vOTU detection in metagenomes compared to viromes (Fig. S3). This suggests that the appropriate breadth thresholds for detection may be different in viromes compared to total metagenomes, with a cut-off ≥75% potentially too conservative for the total metagenomes. However, care should be taken before reducing breadth threshold in other datasets that do not have paired viromes to corroborate viral detection. Regardless, the same pH trends were observed for both viromes and metagenomes.

The distinct soil viromes from the gradient pH 4.5. and 7.5 soils were compared with those recovered from six other soil ecosystems varying in pH, soil type, land use and location and where viral contigs were predicted using the same tools and community standards (Table S4) [2,19,20]. Specifically, using gene-sharing network analysis [20], the number of clusters containing vOTUs from these additional soils and the local gradient (pH 4.5 only, pH 7.5 only or both) were determined (Fig. 2, Table S5). On average, 31% of clusters were shared (range 1-62%) in pairwise comparisons between all soils (Fig. S4).

**Fig. 2:**
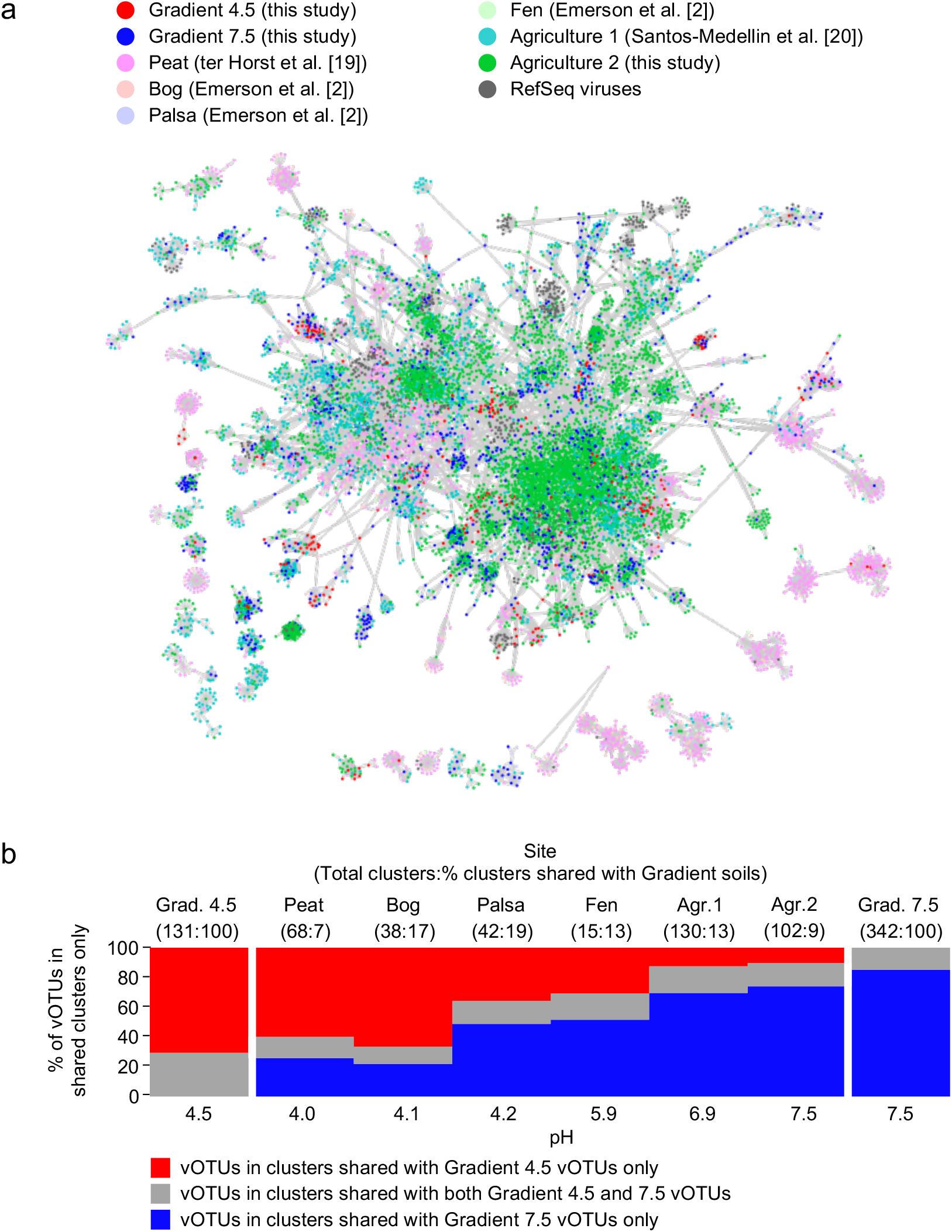
Network analysis describing linkages of Gradient 4.5 and 7.5 vOTUs with six sets of soils samples ranging in pH from 4.0 to 7.5 from Europe and North America (Table S4). a) Gene sharing network of vOTUs showing viral clusters containing ≥25 vOTUs. b) Relative abundance of clusters from each soil that contain vOTUs shared with those from gradient pH 4.5 (blue) and gradient pH 7.5 soil (red) or both (grey). Numbers in parenthesis denote total number and percentage of shared viral clusters with Gradient soils.

A clear trend was observed in comparison with the gradient soils (Fig. 2b) with shared viral clusters in acidic and neutral pH soils more associated with those from the local gradient pH 4.5 or 7.5 soils, respectively.

If individual viruses can infect multiple host populations at different soil pH, there would be a potential for virus community structures to be less distinct over an ecological gradient. However, analysis of samples from a continuous pH gradient demonstrated that, as with prokaryotic community structure, contrasting soil pH results in the selection for virus community structures that are at least as distinct as prokaryote host community structures. Comparison with soil samples of contrasting pH and land-use type also indicated that, as with prokaryote host communities, soil pH correlates with distinct patterns of virus community structures.

## Data availability

Metagenome sequence reads are deposited in NCBI’s GenBank under BioProject accession nos. PRJNA621436–PRJNA621447. Metagenome draft assemblies are accessible through the JGI Genome Portal (DOI: 10.25585/ 1487501). Assembled metagenome-derived 16S rRNA gene sequences are available at ftp://ftp-adn.ec-lyon.fr/. Metagenome sequence reads from ‘Agriculture 2’ site are available through NCBI BioProject PRJNA767554.

## Acknowledgments

This work was funded by an AXA Research Chair awarded to GWN. The sequencing data were generated under JGI Community Science Program proposal 503702 awarded to GWN and CH. The sequencing data from the soil pH gradient was produced by the U.S. Department of Energy Joint Genome Institute, a DOE Office of Science User Facility, which is supported by the Office of Science of the U.S. Department of Energy under Contract No. DE-AC02-05CH11231. The pH gradient experiment is funded through the Scottish Government RESAS 2016-2021 programme. The authors would like to thank Dr. Laurent Pouilloux for assistance with the Newton high performance computing cluster at École Centrale de Lyon. ‘Agriculture 2’ site data collection and analysis was predominantly funded by the UC Davis College of Agricultural and Environmental Sciences and Department of Plant Pathology (new lab start-up to JBE) and also in part by USDA National Institute of Food and Agriculture (NIFA) Hatch project number CA-D-PPA-2464-H and USDA NIFA grant number 2021-67013-34815-0 to JBE. The authors thank Dr. Laura Zinke for collecting and processing the ‘Agriculture 2’ site samples.

## Competing Interests

The authors declare no competing interests.

## 1. Supplementary Methods

### Soil sampling, preparation and analysis of metagenome libraries

Soil was collected from pH 4.5 and 7.5 subplots of a pH gradient that has been maintained since 1961 under an 8-year crop rotation (SRUC, Craibstone Estate, Aberdeen, Scotland; UK grid reference NJ872104) [1]. The soil is a podzol with sandy-loam texture and had supported the growth of potatoes the previous year. Three replicate soil samples were collected from the top 10 cm at 1 m intervals in January 2019. Soil pH (measured in water) were 4.2 ± 0.06 and 7.3 ± 0.04 for the pH 4.5 and 7.5 subplots respectively. Soil was sieved (2 mm mesh size) and stored at 4°C (<1 week) prior to preparing total soil metagenomes and viromes generated as previously described [2]. For non-targeted metagenomes, genomic DNA was extracted from 0.5 g soil using a CTAB buffer phenol:chloroform: isoamyl alcohol bead-beating protocol [3]. DNA from virus particles was extracted with the protocol of Trubl et al. [4] with modifications (see [2]). Library preparation and sequencing was performed at the Joint Genome Institute (JGI) as previously described [2]. De novo contig assembly of 125 to 322 million quality-controlled reads per metagenome (both non-targeted and virome) was performed using MetaSPAdes version 3.13.0 [5]. Taxonomic annotation of metagenomic reads was performed using Kaiju version 1.7.0 [6] with the NCBI nr database (2020-05-25 release) for prokaryote phylum annotation. 16S rRNA gene fragments were extracted and classified from the non-targeted metagenomes using SortMeRNA [7] with the bacterial and archaeal SILVA database. The OTU map files obtained from SortMeRNA were converted to an OTU table using the make_otu_table.py function in QIIME (1.3.0 release) [8]. A heatmap of the relative abundance of OTUs was produced using the heatmaply R package [9] in R v3.6.0

### Virus prediction and community profiling

Contigs of viral origin were predicted from contigs ≥10 kb from both non-targeted and virome metagenomes using VirSorter [10] and DeepVirFinder [11]. For VirSorter analysis, category-1 and −2 free virus categories were retained (representing “most confident” and “likely” virus predictions, respectively) together with category-4 and −5 prophage equivalents. For DeepVirFinder, contigs categorized with a score ≥0.9 and *p*-value ≤0.05 (representing “confident” virus predictions) [11] were retained. Viral contigs were clustered into virus operational taxonomic units (vOTUs) with self-hits of viral contigs from BLASTn analysis (e-value ≤1.0e-50, perc_identity ≥80) removed, and the output file parsed using specific cutoffs (global identity ≥95%, covered length ≥80% of the shorter contig; [12]) by using the command java Parse_BLAST [12]. Single linkage clustering was performed using the command SLC.pl. [12]. To determine the relative abundance of vOTUs, BBMap [13] was used to align reads from each metagenome and virome back to indexed vOTUs with the detection threshold of ≥75% of the contig length, covered ≥1x by reads recruited at ≥90% average nucleotide identity [14]. The relative abundance of each vOTU in each metagenome and virome was calculated based on the length of sequence size and coverage. Abundance was expressed as normalized copies per million reads (CPM). This was calculated as follows: the read per kilobase (RPK) was divided by sequence-length (kb), RPK were summed and divided by 1 million to get a scaling factor, and RPK values were divided by the scaling factor to get the CPM. A heatmap of the relative abundance of vOTUs was produced using the heatmaply R package [9] in R v3.6.0. Taxonomic annotation of viral contigs and vOTUs was performed using vConTACT 2.0 [15] with the RefSeq prokaryote virus database (Release 94; 2019-05-13).

### Virus-host linkage

Three approaches were used to predict virus-host linkages; linking viral protospacer and spacer sequences in clustered regularly interspaced short palindromic repeats (CRISPR) arrays, analysis of shared homologs between host and viral contigs, and gene-sharing network analysis. The CRISPR Recognition Tool (CRT) [16] was used to identify CRISPR arrays from the metagenomes and spacer sequences were extracted using line commands and searched in vOTUs using the Seqkit locate function with 100% sequence identity and positive and negative strand search [17]. However, as only two spacers were identified with exact matches to viral contigs, this approach was not considered further. A homolog-based approach was performed as previously described [18] with a host predicted as the phylum with ≥ 3x shared homologs compared to the second most dominant phylum. Gene-sharing network analysis was conducted using vConTACT 2.0 [15] with the RefSeq prokaryote virus database (Release 94; 2019-05-13).

### Auxiliary metabolic genes

To identify putative virus-encoded auxiliary metabolic genes (AMGs), gene prediction of vOTU contigs was performed using Prodigal v2.6.3 with meta option [19], and putative AMG searches were conducted using VIBRANT [20] and DRAM-v [21], with all vOTUs passed to VirSorter2 using “--prep-for-dramv”, and using a minimum auxiliary score threshold of 3 [22]. AMGs predicted by both tools were manually curated. Putative AMGs at the end of contigs and annotated with multiple functional categories were removed. AMGs predicted using either VIBRANT and DRAM-v were categorized into 14 functional metabolism categories and visualized in a heatmap produced through the heatmaply R package [9] in R v3.6.0.

### Statistical analyses

Unless otherwise noted, all statistical analyses were performed using the vegan package [23] in R v3.6.0. vOTU and 16S rRNA OTU rarefaction and accumulation curves were calculated using the rarecurve and speccacum function respectively. Alpha-diversity indices, Shannon and Simpson indices were calculated using the diversity function, and richness calculated using specnumber. Significant differences in alpha-diversity between pH 4.5 and 7.5 soil was tested using Student’s t-test or Wilcoxon-Mann-Whitney’s test when variances were not homogenous according to Bartlett’s test using the ggpubr R package [24]. Viral and prokaryote community structure of pH 4.5 and 7.5 soil were assessed with principal coordinates analysis using Bray-Curtis dissimilarities calculated using the metaMDS function with normalized vOTU and 16S rRNA OTU relative abundance tables. The impact of soil pH on viral community structure was tested using permutational multivariate analyses of variance (PERMANOVA) on Bray-Curtis dissimilarities with the adonis function.

### Network analysis of soil viral populations from various soil systems

vOTUs from the gradient pH 4.5 and 7.5 soils with those from six other soil systems were used in the network analysis (Table S4). A gene-sharing network of vOTUs was produced using vConTACT 2.0 [15] with the RefSeq prokaryote virus database (Release 94; 2019-05-13).

For ‘Agriculture site 2’ (Table S4), the 7,749 vOTUs were produced as follows: in July and October of 2018, two soil samples were collected from each of 6 one-acre plots. The top 15 cm of soil was collected, sieved through an 8 mm mesh, and kept on ice until storage at −20C. For non-targeted metagenomes (n = 24), genomic DNA was extracted from 0.5 g soil using the PowerSoil Kit (Qiagen, Hilden, Germany), and for viromes (n = 18), virus-like particles were purified from 100 g soil and DNA extracted following the protocol described in Santos-Medellin et al. [25]. Libraries were prepared using the DNA HyperPrep kit (Kapa Biosystems-Roche, Basel, Switzerland) and sequenced on a NovaSeq 6000 at the Technologies and Expression Analysis Core, UC Davis Genome Center. Read quality filtering, contig assembly and clustering, and viral contig identification and vOTU detection was performed as previously described [25].

## 2. Supplementary Tables and Figures

**Table S1:** Number of viral contigs (VC) predicted from total metagenomes (M) and viromes (V) from contigs ≥10 kb using DeepVirFinder only (DVF), VirSorter only (VS), and those predicted by both tools (Both), and resulting number of viral operational taxonomic units (vOTUs).

|  | pH 4.5_1 | pH 4.5_2 | pH 4.5_3 | pH 7.5_1 | pH 7.5_2 | pH 7.5_3 |
| --- | --- | --- | --- | --- | --- | --- |
| Metagenomes | Number of predicted viral contigs |  |  |  |  |  |
| Total contigs | 3193 | 2081 | 3450 | 439 | 399 | 366 |
| DVF (0.9; 0.05) | 22 | 22 | 29 | 2 | 1 | 3 |
| VS (cat1,2,4,5) | 16 | 5 | 20 | 4 | 3 | 2 |
| Both | 8 | 3 | 11 | 1 | 0 | 1 |
| Total M-VC | 30 | 24 | 38 | 5 | 4 | 4 |
| Viromes | Number of predicted viral contigs |  |  |  |  |  |
| Total contigs | 1448 | 1130 | 1166 | 3212 | 2789 | 3788 |
| DVF (0.9; 0.05) | 729 | 611 | 634 | 1746 | 1517 | 2064 |
| VS (cat1,2,4,5) | 81 | 52 | 55 | 148 | 159 | 201 |
| Both | 40 | 27 | 23 | 70 | 65 | 88 |
| Total V-VCs | 770 | 636 | 666 | 1824 | 1611 | 2177 |
| vOTU | Number of vOTU |  |  |  |  |  |
| DVF M-VCs | 5 | 2 | 10 | 1 | 0 | 0 |
| VS M-VCs | 1 | 0 | 3 | 1 | 0 | 1 |
| DVF V-VCs | 161 | 133 | 195 | 368 | 322 | 619 |
| VS V-VCs | 7 | 4 | 11 | 10 | 13 | 43 |
| Total vOTUs | 174 | 139 | 219 | 380 | 335 | 663 |

**Table S2:** Virus-host linkage predicted from gene sharing network and gene homology analysis with viral contigs (VC) and vOTUs.

| Predicted host phylum | Homologue-linked VCs |  | vConTACT 2.0-linked VCs |  | Total linked VCs |  | Homologue-linked vOTUs |  | vConTACT 2.0-linked vOTUs |  | Total linked vOTUs |  |
| --- | --- | --- | --- | --- | --- | --- | --- | --- | --- | --- | --- | --- |
|  | pH 4.5 VCs | pH 7.5 VCs | pH 4.5 VCs | pH 7.5 VCs | pH 4.5 VCs | pH 7.5 VCs | pH 4.5 vOTU | pH 7.5 vOTU | pH 4.5 vOTU | pH 7.5 vOTU | pH 4.5 vOTU | pH 7.5 vOTU |
| Actinobacteria | 464 | 710 | 19 | 19 | 479 (77.6%) | 727 (59.4%) | 135 | 217 | 24 | 37 | 140 (77.7%) | 220 (58.5%) |
| Proteobacteria | 102 | 378 | 0 | 0 | 108 (17.5%) | 387 (31.6%) | 27 | 129 | 9 | 51 | 28 (15.5%) | 129 (34.3%) |
| Acidobacteria | 3 | 1 | 0 | 0 | 3 (0.4%) | 1 (0.08%) | 1 | 0 | 0 | 0 | 1 (0.5%) | 0 (0%) |
| Bacterioidetes | 4 | 62 | 0 | 5 | 4 (0.6%) | 66 (5.3%) | 2 | 17 | 2 | 1 | 4 (2.2%) | 18 (4.7%) |
| Firmicutes | 10 | 3 | 5 | 24 | 15 (2.4%) | 27 (2.2%) | 3 | 2 | 1 | 10 | 4 (2.2%) | 5 (3.3%) |
| Planctomycetes | 1 | 7 | 0 | 0 | 1 (0.1%) | 7 (0.5%) | 1 | 2 | 0 | 0 | 1 (0.5%) | 2 (1.3%) |
| Candidatus Gottesmanbacteria | 0 | 1 | 0 | 0 | 0 (0%) | 1 (0.08%) | 0 | 1 | 0 | 0 | 0 (0%) | 1 (0.2%) |
| Candidatus Moranbacteria | 1 | 0 | 0 | 0 | 1 (0.1%) | 0 (0%) | 0 | 0 | 0 | 0 | 0 (0%) | 0 (0%) |
| Candidatus Schekmanbacteria | 0 | 1 | 0 | 0 | 0 (0%) | 1 (0.08%) | 0 | 0 | 0 | 0 | 0 (0%) | 0 (0%) |
| Candidatus Woesebacteria | 0 | 1 | 0 | 0 | 0 (0%) | 1 (0.08%) | 0 | 0 | 0 | 0 | 0 (0%) | 0 (0%) |
| Chloroflexi | 4 | 3 | 0 | 0 | 4 (0.6%) | 3 (0.2%) | 1 | 0 | 0 | 0 | 1 (0.5%) | 0 (0%) |
| Thaumarchaeota | 2 | 0 | 0 | 0 | 2 (0.3%) | 0 (0%) | 1 | 0 | 0 | 0 | 1 (0.5%) | 0 (0%) |
| Verrucomicrobia | 0 | 2 | 0 | 0 | 0 (0%) | 2 (0.1%) | 0 | 1 | 0 | 0 | 0 (0%) | 1 (0.2%) |

**Table S3:** Alpha-diversity indices calculated from normalized vOTU abundance (from viromes only) and normalized 16S rRNA OTU abundance. Values represent mean ± SD.

|  | Shannon | Simpson | Richness |
| --- | --- | --- | --- |
| pH 4.5 vOTUs | 5.71 ±0.012 | 0.99 ±0.0001 | 942.6 ±22.8 |
| pH 7.5 vOTUs | 6.89 ±0.006 | 0.99 ±0.00005 | 1514.6 ±3.4 |
| pH 4.5 16S OTUs | 6.15 ±0.01 | 0.99 ±0.00025 | 1273 ±62.6 |
| pH 7.5 16S OTUs | 6.58 ±0.03 | 0.99 ±0.00044 | 1519 ±27.0 |

**Table S4:** Details of sites, soils and viral populations (vOTUs) used in network analysis of viral communities.

| Site identity | Location | Coordinates | Soil type | pH | C (%) | N (%) | Number of vOTUs | Reference |
| --- | --- | --- | --- | --- | --- | --- | --- | --- |
| Gradient pH 4.5 | Scotland, UK | 57°11'N,<br>2°12'W | podzol,<br>sandy loam | 4.5 | 6.02 | 0.58 | 532 (this study) | Bartram et al. [26] (see for site description) |
| Gradient pH 7.5 |  |  |  | 7.5 | 7.14 | 0.34 | 1,378 (this study) |  |
| Peat | Minnesota, USA | 18°18'N,<br>65°50'W | peat | 4.0 | 5.80 | 0.95 | 5,006 | ter Horst et al. [27] |
| Bog | Northern Sweden | 68°21'N,<br>19°03'E | peat | 4.1 | 43.18 | 0.80 | 703 | Emerson et al. [28] |
| Palsa |  |  | peat | 4.2 | 35.07 | 1.17 | 794 |  |
| Fen |  |  | peat | 5.9 | 37.59 | 1.34 | 334 |  |
| Agriculture 1 | California, USA | 38°32'08"N,<br>121°46'22"W | Yolo silt<br>loam | 6.9 | 1.10 | 0.12 | 4,065 | Santos-Medellin et al. [25] |
| Agriculture 2 | California, USA | 38°32'42"N,<br>121°52'36" W | Rincon silty<br>clay loam | 7.5 | 1.36 | 0.15 | 7,749 (this study) | Wolf et al. [29] (see for site description) |

**Table S5:** Summary of gene sharing network analysis and the number of vOTUs in clusters from various soils shared with those from gradient pH 4.5 and 7.5 soils.

| Site identity | Number of nodes (vOTUs) | Number of singleton clusters | Number of vOTUs clustered with unique gradient pH 4.5 vOTUs | Number of vOTUs clustered with unique gradient pH 7.5 vOTUs | Number of vOTUs clustered with both gradient pH 4.5 and 7.5 vOTUs |
| --- | --- | --- | --- | --- | --- |
| Gradient pH 4.5 | 467 | 9 | 189 | 0 | 84 |
| Gradient pH 7.5 | 1,272 | 24 | 0 | 590 | 105 |
| Peat | 4,456 | 95 | 135 | 61 | 32 |
| Bog | 646 | 4 | 46 | 16 | 8 |
| Palsa | 640 | 16 | 25 | 35 | 11 |
| Fen | 265 | 3 | 7 | 12 | 4 |
| Agriculture 1 | 3,843 | 44 | 48 | 272 | 69 |
| Agriculture 2 | 7,282 | 147 | 96 | 708 | 150 |
| RefSeq | 2,568 | 101 | 9 | 28 | 8 |
| Total | 18,871 | 443 |  |  |  |

**Fig. S1:**
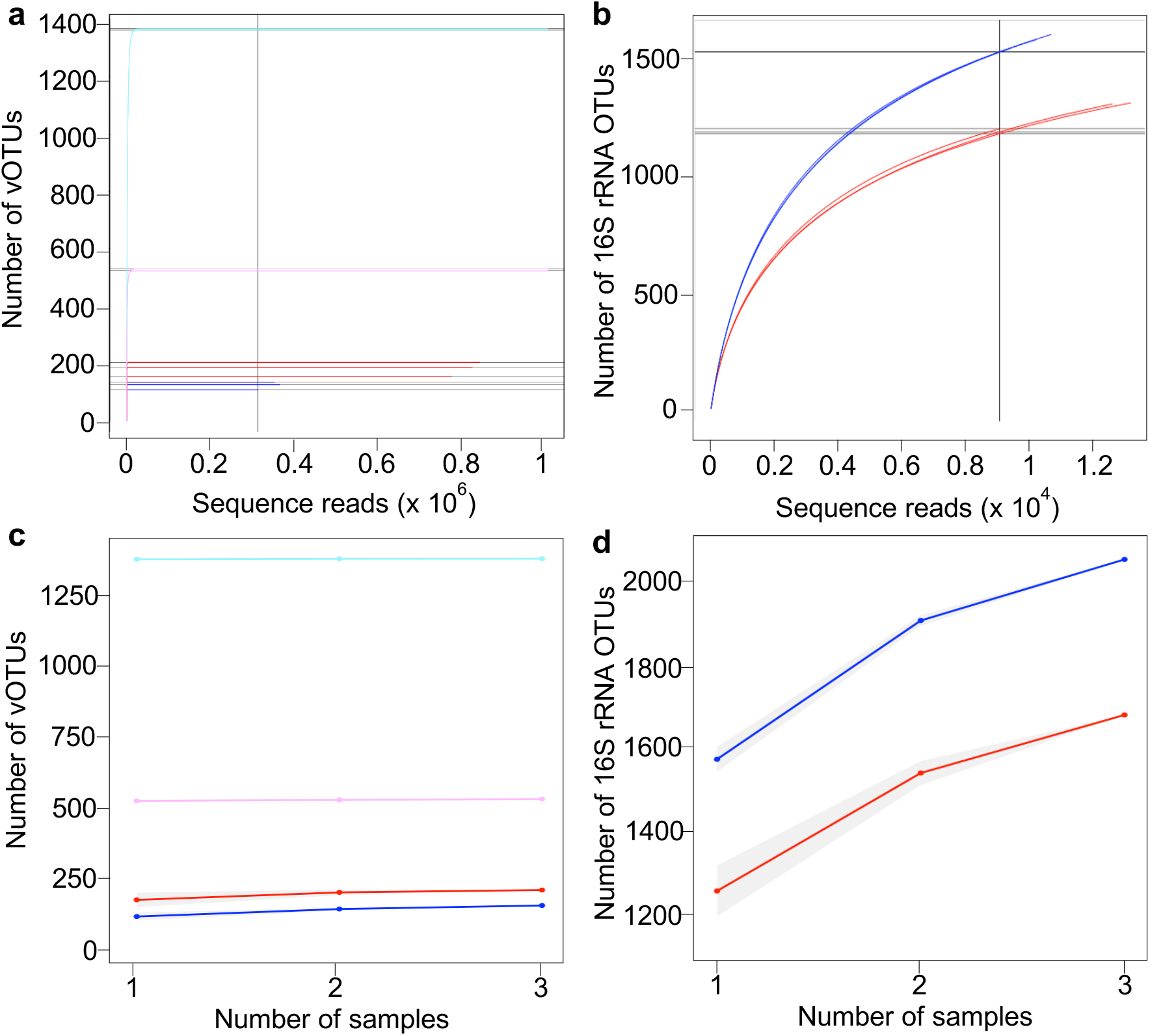
Rarefaction and accumulation curves for vOTUs in total metagenomes (pH 4.5, red; pH 7.5, blue; n=3) and viromes (pH 4.5, pink; pH 7.5, cyan; n=3) a) and c), and 16S rRNA OTUs in total metagenomes b) and d).

**Fig. S2:**
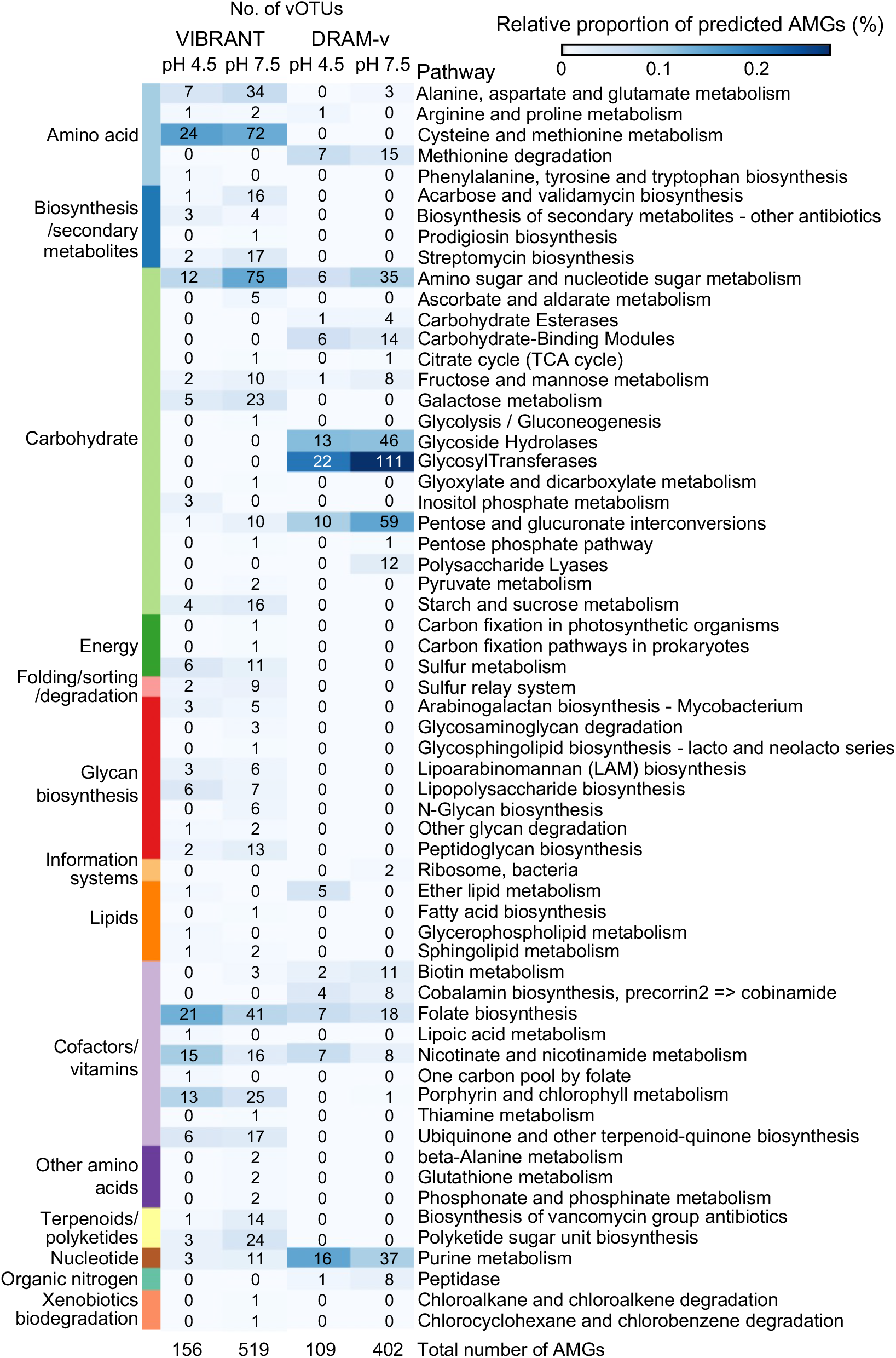
Metabolic category and function of auxiliary metabolic genes (AMGs) in vOTUs from gradient 4.5 and 7.5 soil. The number and relative proportion (based on annotated genes) are given for both VIBRANT and DRAM-v.

**Fig. S3:**
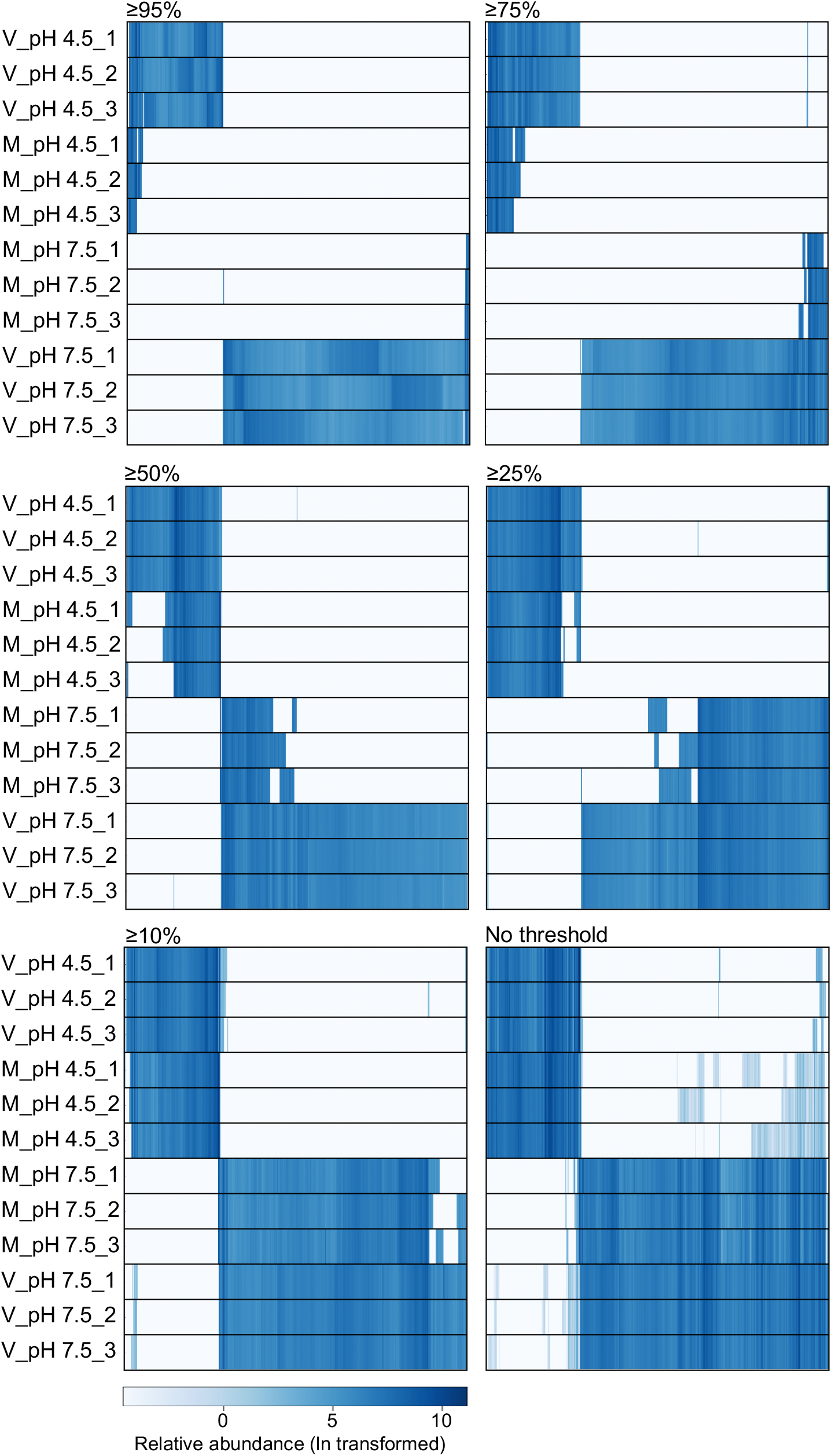
Effect of breadth (length of contig covered by mapped reads) thresholds for defining detection of vOTU contigs. Heatmaps present the normalized relative abundance of individual vOTUs in gradient pH 4.5 and 7.5 soil viromes (V) and metagenomes (M).

**Fig. S4:**
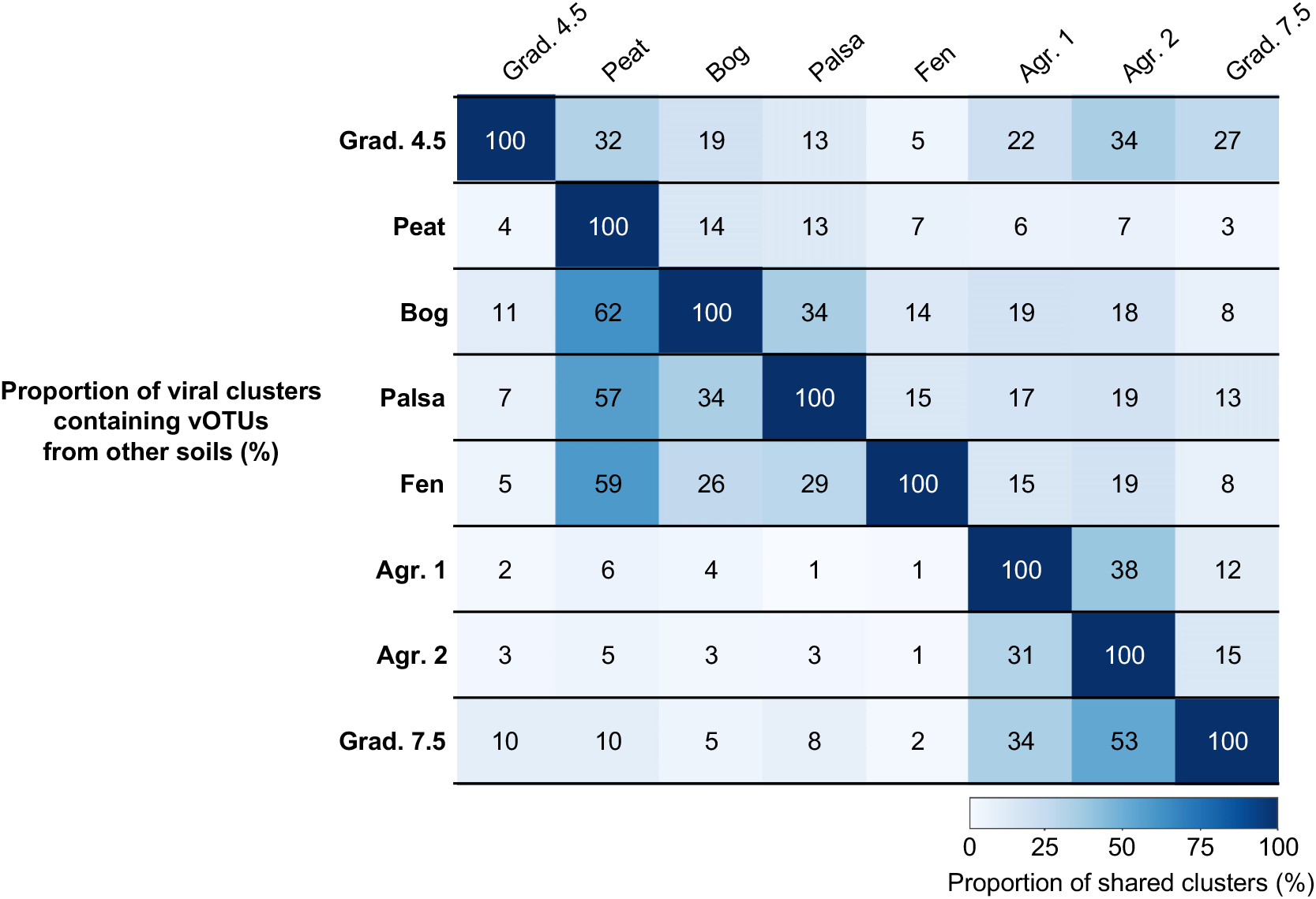
Percentage of viral clusters shared between soils based on the gene sharing network analysis.

## Notes

### Competing Interest Statement

The authors have declared no competing interest.

## References

1. Pratama AA, van Elsas JD. The ‘neglected’ soil virome – potential role and impact. Trends Microbiol. 2018;26;649–662.

2. Emerson JB, Roux S, Brum JR, Bolduc B, Woodcroft BJ, Jang HB, et al. Host-linked soil viral ecology along a permafrost thaw gradient. Nat Microbiol. 2018;3:870–80.

3. Williamson KE, Fuhrmann JJ, Wommack KE, Radosevich M. Viruses in Soil Ecosystems: An Unknown Quantity Within an Unexplored Territory. Annu Rev Virol. 2017;4:201–219.

4. Kuzyakov and Mason-Jones. Viruses in soil: Nano-scale undead drivers of microbial life, biogeochemical turnover and ecosystem functions. Soil Biol Biochem. 2018;127:305–317.

5. Trubl G, Jang HB, Roux S, Emerson JB, Solonenko N, Vik DR, et al. Soil viruses are underexplored players in ecosystem carbon processing. mSystems. 2018;3.

6. Vos M, Birkett PJ, Brich E, Griffiths RJ, Buckling A. Local adaptation of bacteriophages to their bacterial hosts in soil. Science. 2009;14:833.

7. Srinivasiah S, Lovett J, Ghosh D, Roy K, Fuhrmann JJ, Radosevich M, et al. Dynamics of autochthonous soil viral communities parallels dynamics of host communities under nutrient stimulation. FEMS Microbiol Ecol. 2015;91:fiv063.

8. de Jonge PA, Nobrega FL, Brouns SJJ, Dutilh BE. Molecular and evolutionary determinants of bacteriophage host range. Trends Microbiol. 2019;27:51–63.

9. Trubl G, Solonenko N, Chittick L, Solonenko SA, Rich VI, Sullivan MB. Optimization of viral suspension methods for carbon-rich soils along a permafrost thaw gradient. PeerJ. 2016;17:e1999.

10. Lammel DR, Barth G, Ovaskainen O, Cruz LM, Zanatta JA, Ryo M, et al. Direct and indirect effects of a pH gradient bring insights into the mechanisms driving prokaryotic community structures. Microbiome. 2018;6:106.

11. Bahram M, Hildebrand F, Forslund SK, Anderson JL, Soudzilovskaia HA, Bodegom PM, et al. Structure and function of the global topsoil microbiome. Nature. 2018;560:233–237.

12. Lauber CL, Hamady M, Knight R, Fierer N. Pyrosequencing-based assessment of soil pH as a predictor of soil bacterial community structure at the continental scale. Appl Environ Microbiol. 2009;75:5111–5120.

13. Nicol GW, Leininger S, Schleper C, Prosser JI. The influence of soil pH on the diversity, abundance and transcriptional activity of ammonia oxidizing archaea and bacteria. Environ Microbiol. 2008;10:2966–78.

14. Bartram AK, Jiang X, Lynch MD, Masella AP, Nicol GW, Dushoff J, et al. Exploring links between pH and bacterial community composition in soils from the Craibstone Experimental Farm. FEMS Microbiol Ecol. 2014;87:403–15.

15. Lee S, Sieradzki ET, Nicolas AM, Walker RL, Firestone MK, et al. Methane-derived carbon flows into host-virus networks at different trophic levels in soil. PNAS. 2021;10:e2105124118.

16. Jang HB, Bolduc B, Zablocki O, Kuhn JH, Roux S, Adriaenssens EM, et al. Taxonomic assignment of uncultivated prokaryotic virus genomes is enabled by gene-sharing networks. Nat. Biotechnol. 2019;37:632–639.

17. Al-Shayeb B, Sachdeva R, Chen LX, Ward F, Munk P, Devoto A, et al. Clades of huge phages from across Earth’s ecosystems. Nature. 2020;578:425–431.

18. Sobsey MD, Dean CH, Knuckles ME, Wagner RA. Interactions and survival of enteric viruses in soil material. Appl Environ Microbiol. 1980;40:92–101.

19. ter Horst AM, Santos-Medellin C, Sorensen JW, Zinke LA, Wilson RM, Johnston ER, et al. Minnesota peat viromes reveal terrestrial and aquatic niche partitioning for local and global viral populations. Microbiome (in press).

20. Santos-Medellin C, Zinke LA, ter Horst AM, Gelardi DL, Parikh SJ, Emerson JB. Viromes outperform total metagenomes in revealing the spatiotemporal patterns of agricultural soil viral communities. ISME J. 2021;15:1956–1970.

## Supplementary References

1. Kemp JS, Paterson E, Gammack SM, Cresser MS, Killham K. Leaching of genetically modified Pseudomonas fluorescens through organic soils: influence of temperature, soil pH, and roots. Biol. Fert. Soils. 1992;13:218–224.

2. Lee S, Sieradzki ET, Nicolas AM, Walker RL, Firestone MK, et al. Methane-derived carbon flows into host-virus networks at different trophic levels in soil. PNAS USA. 2021;10:e2105124118.

3. Nicol GW, Prosser JI. Strategies to determine diversity, growth and activity of ammonia oxidising archaea in soil. Meth. Enzymol. 2011;496:3–34.

4. Trubl G, Solonenko N, Chittick L, Solonenko SA, Rich VI, Sullivan MB. Optimization of viral suspension methods for carbon-rich soils along a permafrost thaw gradient. PeerJ. 2016;17:e1999.

5. Nurk S, Meleshko D, Korobeynikov A, Pevzner PA. metaSPAdes: a new versatile metagenomic assembler. Genome Res. 2017;27:824–834.

6. Menzel P, Ng KL, Krogh A. Fast and sensitive taxonomic classification for metagenomics with Kaiju. Nat. Commun. 2016;7:11257.

7. Kopylova E, Noé L, Touzet H. SortMeRNA: fast and accurate filtering of ribosomal RNAs in metatranscriptomic data. Bioinformatics 2012;28:3211–3217.

8. Caporaso JG, Kuczynski J, Stombaugh J, Bittinger K, Bushman FD, Cpstello EK et al. QIIME allows analysis of high-throughput community sequencing data. Nat. Methods 2010;7:335–336.

9. Galili T, O’Callaghan A, Sidi J, Sievert C. heatmaply: an R package for creating interactive cluster heatmaps for online publishing. Bioinformatics. 2018;34:1600–1602.

10. Roux S, Enault F, Hurwitz BL, Sullivan MB. VirSorter: mining viral signal from microbial genomic data. PeerJ. 2015;3:e985.

11. Ren J, Song K, Deng C, Ahlgren NA, Fuhrman JA, Li Y, Xie X, Poplin R, Sun F. Identifying viruses from metagenomic data using deep learning. Quant. Biol. 2020;8:64–77.

12. Paez-Espino D, Pavlopoulos GA, Ivanova NN, Kyrpides NC. Nontargeted virus sequence discovery pipeline and virus clustering for metagenomic data. Nat. Protoc. 2017;12:1673–1682.

13. Bushnell B. BBTools software package. 2014. http://sourceforge.net/projects/bbmap.

14. Roux S, Adriaenssens EM, Dutilh BE, Koonin EV, Kropinski AM, Krupovic M, et al. Minimum Information about an Uncultivated Virus Genome (MIUViG). Nat. Biotechnol. 2019;37:29–37.

15. Jang HB, Bolduc B, Zablocki O, Kuhn JH, Roux S, Adriaenssens EM, et al. Taxonomic assignment of uncultivated prokaryotic virus genomes is enabled by gene-sharing networks. Nat. Biotechnol. 2019;37:632–639.

16. Bland C, Ramsey TL, Sabree F, Lowe M, Brown K, Kyrpides NC, et al. CRISPR Recognition Tool (CRT): a tool for automatic detection of clustered regularly interspaced palindromic repeats. BMC Bioinformatics. 2007;8:209.

17. Shen W, Le S, Li Y, Hu F. SeqKit: A Cross-Platform and Ultrafast Toolkit for FASTA/Q File Manipulation. PLoS One. 2016;11:e0163962.

18. Al-Shayeb B, Sachdeva R, Chen LX, Ward F, Munk P, Devoto A, et al. Clades of huge phages from across Earth’s ecosystems. Nature. 2020;578:425–431.

19. Hyatt D, Chen GL, Locascio PF, Land ML, Larimer FW, Hauser LJ. Prodigal: prokaryotic gene recognition and translation initiation site identification. BMC Bioinformatics. 2010;11:119.

20. Kieft K, Zhou Z, Anantharaman K. VIBRANT: automated recovery, annotation and curation of microbial viruses, and evaluation of viral community function from genomic sequences. Microbiome. 2020;8:90.

21. Shaffer M, Borton MA, McGivern BB, Zayed AA, La Rosa SL, Solden LM, et al. DRAM for distilling microbial metabolism to automate the curation of microbiome function. Nucleic Acids Res. 2020;48:8883–8900.

22. Guo J, Bolduc B, Zayed AA, Varsani A, Dominguez-Huerta G, Delmont TO, et al. VirSorter2: a multi-classifier, expert-guided approach to detect diverse DNA and RNA viruses. Microbiome. 2021;9:37.

23. Oksanen JF, Blanchet G, Friendly M, Kindt R, Legendre P, McGlinn D, et al. vegan:Community Ecology Package. R package version 2.5-6. 2019. https://CRAN.R-project.org/package=vegan

24. Kassambara A. ggpubr: ‘ggplot2’ Based Publication Ready Plots. R package version 0.3.0. 2020. https://CRAN.R-project.org/package=ggpubr

25. Santos-Medellin C, Zinke LA, ter Horst AM, Gelardi DL, Parikh SJ, Emerson JB. Viromes outperform total metagenomes in revealing the spatiotemporal patterns of agricultural soil viral communities. ISME J. 2021;15:1956–1970.

26. Bartram AK, Jiang X, Lynch MD, Masella AP, Nicol GW, Dushoff J, et al. Exploring links between pH and bacterial community composition in soils from the Craibstone Experimental Farm. FEMS Microbiol. Ecol. 2014;87:403–15.

27. ter Horst AM, Santos-Medellin C, Sorensen JW, Zinke LA, Wilson RM, Johnston ER, et al. Minnesota peat viromes reveal terrestrial and aquatic niche partitioning for local and global viral populations. Microbiome (in press).

28. Emerson JB, Roux S, Brum JR, Bolduc B, Woodcroft BJ, Jang HB, et al. Host-linked soil viral ecology along a permafrost thaw gradient. Nat Microbiol. 2018;3:870–880.

29. Wolf KM, Torbert EE, Bryant D, Burger M, Denison RF, Herrera et al. The century experiment: the first twenty years of UC Davis’ Mediterranean agroecological experiment. Ecology 2018;99:503.

